# Dynamic local mRNA distribution and translation influence the postnatal molecular maturation of perivascular astrocytic processes

**DOI:** 10.1101/2023.07.25.550497

**Authors:** Katia Avila-Gutierrez, Leila Slaoui, Rodrigo Alvear, Esther Kozlowski, Marc Oudart, Emma Augustin, Philippe Mailly, Héloïse Monnet, Virginie Mignon, Bruno Saubaméa, Anne-Cécile Boulay, Martine Cohen-Salmon

## Abstract

Astrocytes (the main glial cells in the brain) are highly ramified and send out perivascular processes (PvAPs) that entirely sheathe the brain’s blood vessels. PvAPs are equipped with an enriched molecular repertoire that sustains astrocytic regulatory functions at the vascular interface. In the mouse, PvAP development starts after birth and is essentially complete by postnatal day (P) 15. Progressive molecular maturation also occurs over this period, with the acquisition of proteins enriched in PvAPs. The mechanisms controlling the development and molecular maturation of PvAPs have not been extensively characterized. We reported previously that mRNAs are distributed unequally in mature PvAPs and are locally translated. Since dynamic mRNA distribution and local translation influence the cell’s polarity, we hypothesized that they might sustain the postnatal maturation of PvAPs. Here, we used a combination of molecular biology and imaging approaches to demonstrate that the development of PvAPs is accompanied by the transport of mRNA and polysomal mRNA into PvAPs, the development of a rough endoplasmic reticulum (RER) network and Golgi cisternae, and local translation. By focusing on genes and proteins that are selectively or specifically expressed in astrocytes, we characterized the developmental profile of mRNAs, polysomal mRNAs and proteins in PvAPs from P5 to P60. Furthermore, we found that distribution of mRNAs in PvAPs is perturbed in a mouse model of megalencephalic leukoencephalopathy with subcortical cysts. Lastly, we found that some polysomal mRNAs polarized progressively towards the PvAPs. Our results indicate that dynamic mRNA distribution and local translation influence the postnatal maturation of PvAPs.

**Summary statement:** Local translation operates during the postnatal development of perivascular astrocyte processes and might contribute to their molecular maturation.

## Introduction

Astrocytes are abundant glial cells in the brain.They influence the cerebrovascular system through regulation of the blood-brain barrier (BBB), endothelial transport, the recruitment of immune cells by endothelial cells, perivascular homeostasis, blood flow and metabolite transfer to neurons (Cohen-Salmon et al., 2021). Most of these regulatory actions are exerted at a specific interface between the astrocyte and the brain vessel, referred to as the gliovascular unit. Astrocytes are highly ramified and have perivascular astrocytic processes (PvAPs, also known as endfeet) that fully cover the brain’s blood vessels. Recent research results suggest that all cortical astrocytes have PvAPs and contact between 1 to 3 vessels (Hosli et al., 2022). The gliovascular unit develops after birth and requires the densification of the vascular network (Coelho-Santos et al., 2021; Coelho-Santos and Shih, 2020), the maturation of vascular smooth muscle cell contractility (Slaoui et al., 2023), the recruitment of perivascular macrophages (Karam et al., 2022) and fibroblasts (Jones et al., 2023), and the production and maturation of astrocytes and PvAPs. In the cortex, astrocytogenesis starts at around the time of birth and peaks during the following week. Astrocytogenesis is followed by a phase during which the astrocytes ramify progressively and acquire their final shape (Clavreul et al., 2019). We recently reported that (i) PvAPs in the mouse cortex and hippocampus are formed around postnatal (P) day 5, and (ii) astrocytic perivascular coverage increases rapidly between P5 and P10 and is almost complete by P15 (Gilbert et al., 2021). Postnatally, the PvAPs also go through a molecular maturation phase, with the progressive acquisition of a specific molecular repertoire that sustains the cells’ regulatory functions at the vascular interface. The best-characterized PvAP proteins include the water channel aquaporin 4 (Aqp4) and the inward rectifying potassium channel Kir4.1, which controls perivascular water and potassium homeostasis (Lunde et al., 2015). Aqp4 is also involved in intrapenchymal drainage mechanisms (Iliff et al., 2012). Other well-characterized PvAP proteins include the glutamate transporter Glt1 (which controls perivascular glutamate homeostasis), the gap junction proteins connexin (Cx)30 and Cx43 (which regulate BBB integrity and immune system homeostasis) (Boulay et al., 2015a; Ezan et al., 2012), MLC1, and GlialCAM. The latter two proteins form an adhesion complex between astrocyte PvAPs that controls the development of the astrocyte’s perivascular coverage and influences arteriolar vascular smooth muscle cell contractile properties, cerebral blood flow, neurovascular coupling, and drainage of interstitial fluid in the brain (Gilbert et al., 2021; Gilbert et al., 2019).

The acquisition and development of the PvAPs’ specific molecular repertoire have not been extensively characterized. We have previously reported that local translation occurs in PvAPs (Boulay et al., 2017) and that the polysomes in PvAPs contain mRNAs coding for the key proteins mentioned in the previous paragraph (Boulay et al., 2017). Dynamic mRNA distribution and local translation are common in cells with a complex morphology. These local processes mediate cell polarity and allow cells to adapt to local stimuli without being dependent on somatic protein synthesis, which is sometimes far removed from the regulatory site. In the brain, local translation mechanisms have been extensively studied in neurons (Ament and Poulopoulos, 2023) but have also been evidenced in oligodendrocytes, astrocytes (reviewed in (Blanco-Urrejola et al., 2021; Mazare et al., 2021; Meservey et al., 2021) and (most recently) microglia (Vasek et al., 2023)).

In the present study, we hypothesized that dynamic mRNA distribution and local translation sustain the postnatal maturation of PvAPs. We therefore studied the subcellular organization of developing PvAPs in mice and the distribution of mRNAs, polysomal mRNAs and protein levels in a normal (healthy) context and in a mouse model of megalencephalic leukoencephalopathy with subcortical cysts (MLC). Furthermore, we studied the polarity of polysomal mRNAs in astrocytes and PvAPs. Our results show that mRNA distribution and local translation occur in developing PvAPs and might contribute to the acquisition of the processes’ molecular identity.

## Results

### Translation occurs in newly formed PvAPs

We first sought to determine whether differential mRNA distribution and local translation occur in newly formed PvAPs. By using fluorescent *in situ* hybridization (FISH) to look for mRNAs in PvAPs on P5 dorsal cortical sections, we detected the astrocyte-specific *Aqp4* and *Fabp7* mRNAs (**Fig. 1A**). We used isolectin B4 (IB4) to visualize the brain’s blood vessels. The FISH dots were detected in the astrocyte’s soma and processes and in the PvAPs surrounding the IB4-labeled blood vessels (**Fig. 1A**). We found that in the PvAPs on P5, *Fapb7* mRNA was more abundant than *Aqp4* mRNA (**Fig. 1B**).

**Figure 1:**
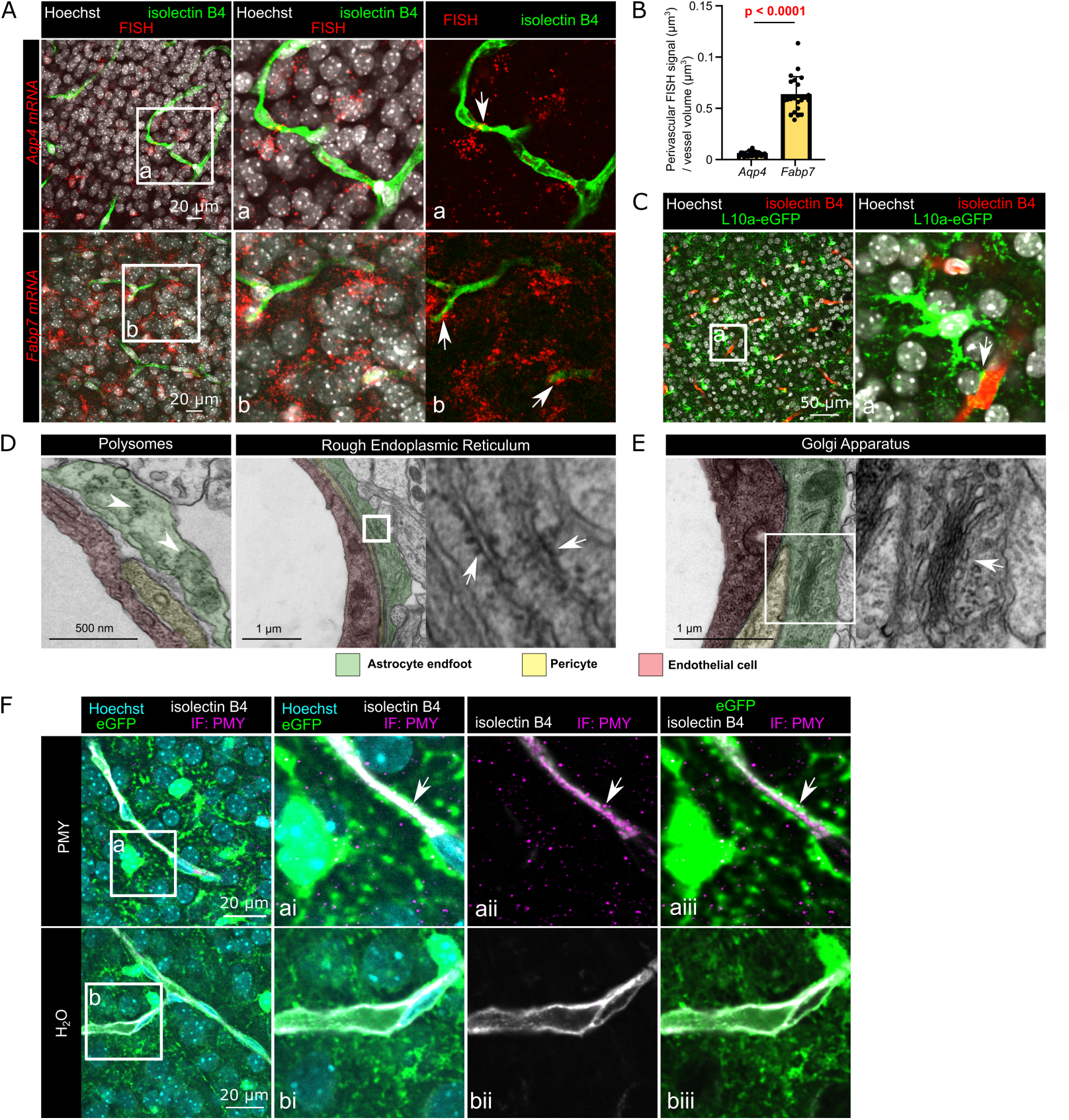
Translation occurs in newly formed PvAPs. **A.** Representative confocal microscopy images of the FISH detection of *Aqp4* and *Fabp7* mRNAs in the cortex on P5. Vessels (in green) were stained with isolectin B4. Nuclei (in white) were stained with Hoechst reagent. Boxed areas are enlarged in the images **(a)** and **(b)**. White arrows indicate FISH dots at the surface of microvessels. **B.** The perivascular FISH signal, normalized against the vascular volume on P5. A histogram of the data shows the mean ± SD (*Aqp4*: n=32 images in three mice; *Fabp7*: n=22 images in two mice); two-tailed Mann-Whitney test. The raw data are presented in **Table S2**. **C.** Representative confocal microscopy images of the immunofluorescence detection of astrocytic ribosomes tagged with eGFP in the Aldhl1:L10a-eGFP mouse cortex on P5. Vessels were stained with isolectin B4 (in red). Nuclei (in white) were stained with Hoechst reagent. The boxed area is enlarged in the image (**a**). A white arrow indicates L10a-eGFP immunofluorescence in PvAPs. **D. E.** Representative TEM images of the gliovascular unit in the cortex on P5. **D.** Images are presented in artificial colors, with PvAPs in green, endothelial cells in pink, and pericytes in yellow. Boxed areas in the PvAP are enlarged in the right-hand images. White arrowheads indicate polysomes. White arrows indicate RER. **E.** PvAP with the Golgi apparatus (white arrow). The boxed area in the PvAP is enlarged in the right-hand image. **F.** Representative confocal microscopy images in the cortex of Aldh1l1:eGFP mice injected with puromycin (PMY) or water on P5. Puromycilated nascent protein chains are labeled in pink. Vessels (in white) were stained with isolectin B4. Nuclei (in cyan) were stained with Hoechst reagent. (**a, b**) boxed areas are enlarged in the right-hand images showing the merge for all labelings (**ai, bi**), isolectin B4 and PMY (**aii, bii**), eGFP, isolectin B4 and PMY (**aiii, biii**). White arrows indicate PMY dots colocalizing with the perivascular eGFP fluorescence.

We next checked for the presence of ribosomes in P5 PvAPs. First, we studied the Aldh1l1:L10a-eGFP transgenic mouse line, which expresses the eGFP-tagged ribosomal protein Rpl10a specifically in astrocytes (Heiman et al., 2008). On P5 cortical sections, we found eGFP-labeled ribosomes both in the astrocyte soma and PvAPs (**Fig. 1C**). The presence of ribosomes in PvAPs on P5 was further confirmed by transmission electron microscopy (TEM) (**Fig. 1D**). We focused on PvAPs surrounding 5- to 20-µm-diameter vessels in mouse cortices. Remarkably, we observed both free polysomal structures and elongated endoplasmic reticulum (ER) membranes covered with ribosomes (i.e. rough ER (RER)) (**Fig. 1D**). We also observed Golgi cisternae in some of the PvAPs (**Fig. 1E**). The presence of mRNAs, polysomes and organelles for protein translation and maturation suggested that translation occurs in PvAPs on P5. To test this hypothesis, we performed puromycilation experiments (**Fig. 1F**). Puromycin (PMY) is an aminoglycoside antibiotic that mimics charged tRNA-Tyr and is incorporated into the ribosome A site. It induces premature translation termination by ribosome-catalyzed covalent incorporation into the nascent carboxy-terminal chain; hence, immunodetection of PMY *in situ* allows the detection of translation events (Pestka and Brot, 1971). Here, P5 Aldh1l1:eGFP transgenic mice (in which the astrocyte’s cytoplasm is filled with eGFP (Srinivasan et al., 2016)) were exposed to intraperitoneally injected PMY, which was then detected on brain sections by immunofluorescence (**Fig. 1F**). Again, brain vessels were labelled using IB4. We detected a PMY signal co-localized with eGFP and proximal to the IB4 staining, which suggested the presence of translation events in PvAPs (**Fig. 1F**).

Taken as a whole, these results showed that mRNA distribution and local translation take place in newly formed PvAPs and that the latter are equipped with organelles dedicated to translation and protein maturation.

### The subcellular organization of translation in developing PvAPs

As mentioned in the introduction, astrocyte perivascular coverage develops postnatally until P15 (Gilbert et al., 2021) and PvAPs follow a molecular maturation process (Gilbert et al., 2019; Lunde et al., 2015). Here, we sought to determine whether the subcellular organization of translation in PvAPs also changed postnatally. Firstly, we focused on the ribosomes by leveraging the features of the Aldh1l1:L10a-eGFP transgenic mouse line. We quantified the volume of perivascular Rpl10a-eGFP immunofluorescence from P5 to P15 in the cortex and again used IB4 labelling to image the brain vessel surface. The volume rose progressively from P5 to P60 (**Fig. 2A, B**). The same progression was also observed for cytoplasmic perivascular eGFP on Aldh1l1:eGFP cortical sections (Srinivasan et al., 2016) **(Fig. 2C, D**). We next used TEM to study the ultrastructure of PvAPs surrounding 5- to 20-µm-diameter vessels in the cortex on P5 **(Fig. 1D, E)**, P10, P15, and P60 **(Fig. 2E)**. The proportion of the PvAP area containing ribosomes, polysomes and/or RER was around 50% and did not change during development (**Fig. 2F**). Likewise, the length of the RER (normalized against the PvAP coverage) did not change significantly (**Fig. 2G**).

**Figure 2:**
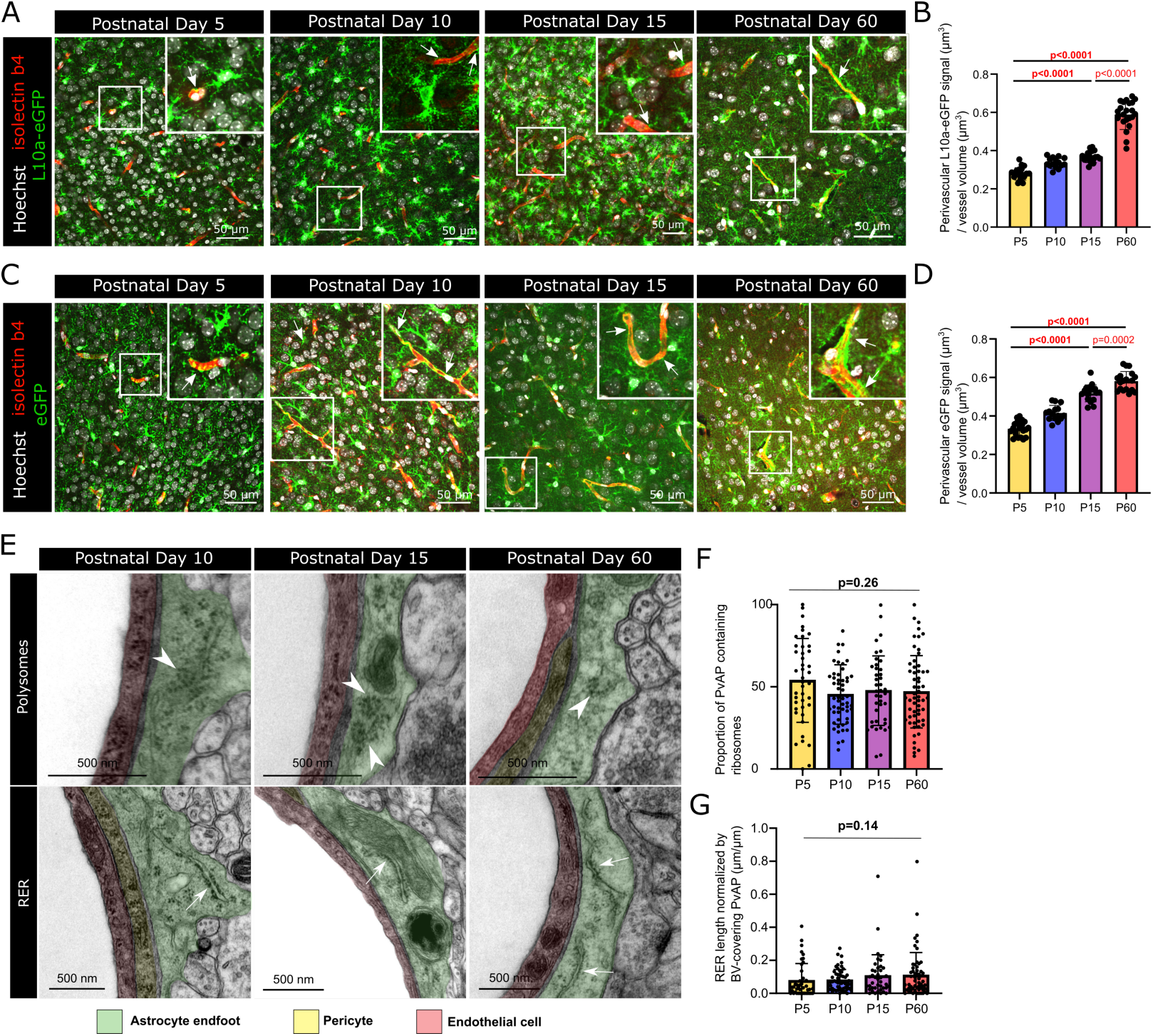
The subcellular organization of translation in developing PvAPs. **A.** Representative confocal microscopy images of the immunofluorescent detection of astrocytic eGFP-tagged ribosomes from Aldh1l1:L10a-eGFP mouse cortex on P5, P10, P15, and P60. Blood vessels (in red) were stained with isolectin B4. Nuclei (in white) were stained with Hoechst reagent. Boxed areas are enlarged in the same image. White arrows indicate L10a-eGFP perivascular fluorescence. **B.** Quantification of the volume of Aldh1l:L10a-eGFP perivascular fluorescence on P5, P10, P15, and P60. Perivascular cytosolic L10a-eGFP fluorescence was normalized against the vessel volume. The data are presented as the mean ± SD (N = three mice per stage, 17 images on P5, 15 on P10, 18 on P15, 23 on P60)); A one-way ANOVA was followed by Tukey’s test for multiple comparisons. **C.** Representative confocal microscopy images of the fluorescence detection of eGFP in astrocytic cytoplasm in the Aldh1l1:eGFP mouse cortex on P5, P10, P15, and P60. Vessels (in red) were stained with isolectin B4. Nuclei (in white) were stained with Hoechst reagent. **D.** Quantification of Aldh1l1:eGFP perivascular fluorescence on P5, P10, P15, and P60. The perivascular cytosolic eGFP fluorescence was normalized against the vessel volume. The data are quoted as the mean ± SD. A one-way ANOVA was followed by Tukey’s test for multiple comparisons (N= three mice per stage; 26 images on P5, 15 on P10, 18 on P15, and 18 on P60). **E.** Representative TEM images of the gliovascular unit in the cortex on P10, P15, and P60, showing the presence of polysomes (white arrowheads) and RER (white arrows) in PvAPs (the images on P5 are shown in Fig. 1D). Images are presented in artificial colors, with perivascular astrocytic process in green, endothelial cells in pink, and pericytes in yellow. **F, G.** Quantification of the proportion of PvAPs containing polysomes and/or RER on TEM images (**F**) and the length of RER normalized against the PvAP coverage (**G**). The data are plotted as the mean ± SD. A one-way ANOVA was followed by Tukey’s test for multiple comparisons (**F**), or a Kruskal-Wallis test was followed by Dunn’s test for multiple comparisons (**G**) (mice per sample: 4 for P5; 3 for P10; 3 for P15; 4 for P60). The raw data are presented in **Table S2**.

Taken as a whole, these observations suggest that subcellular translation machinery is already present in PvAPs at P5. The amount of perivascular cytoplasmic eGFP showing the progressive postnatal development of PvAPs and the amount of eGFP-tagged ribosomes in PvAPs are correlated, indicating that the formation of PvAPs is accompanied by the migration of ribosomes. However, not all PvAPs contains RER and/or polysomes and the proportion of PvAPs containing RER and/or polysomes does not change significantly during postnatal development.

### Analysis of mRNAs in PvAPs during postnatal development

We next sought to determine which mRNAs were distributed across PvAPs during postnatal development. We had previously reported that mRNA-containing PvAPs remain attached to mechanically purified brain microvessels (MVs) (Boulay et al., 2017; Boulay et al., 2015b). Furthermore, we also recently described the transcriptome of mechanically purified dorsal cortex MVs on P5 and P15 (Slaoui et al., 2023) (**Fig. 3A**). In the present study, we leveraged our knowledge of this transcriptome to detect changes over time in MV’s mRNAs that are selectively or specifically expressed in astrocytes (and thus potentially present in PvAPs), according to recently published single-cell RNA sequencing (RNASeq) datasets for adult brain vascular cells (He et al., 2016; Vanlandewijck et al., 2018). A transcript was selected when it was (i) detected in more than 60% of the astrocytes, (ii) detected in less than 12% of other vascular and neural cells, and (iii) expressed at a higher level in astrocytes than in other cells (Log_2_FC ≥ 1.5) (**Table S1**) (**Fig. 3A**). This approach enabled us to identify 140 transcripts selectively or specifically expressed in astrocytes (**Table S1**). Next, we analyzed the expression of each transcript in our cortical MV transcriptome on P5 and P15 (**Table S1**). Of the 140 transcripts, 77 were detected in PvAPs on P5 (reads per kilobase per million mapped reads (RPKM)_P5_ ≥ 10) and 84 were detected on P15 (RPKMP_15_ ≥ 10). Seventy transcripts had similar levels in PvAPs on P5 and on P15 (RPKM_P5_ ≥ 10 or RPKMP_15_ ≥ 10 and -1 < log_2_FC <1 or padj > 0.05) and 14 transcripts were upregulated on P15 vs. P5 (log_2_FC ≥ 1, RPKMP_15_ ≥ 10 and padj ≤ 0.05). Ten of the 14 upregulated transcripts were detected on P15 only (RPKM < 10 on P5). Eight transcripts were downregulated on P15, relative to P5 (log_2_FC ≤ -1 and padj ≤ 0.05) (**Fig. 3B, C**; **Table S1**).

**Figure 3.**
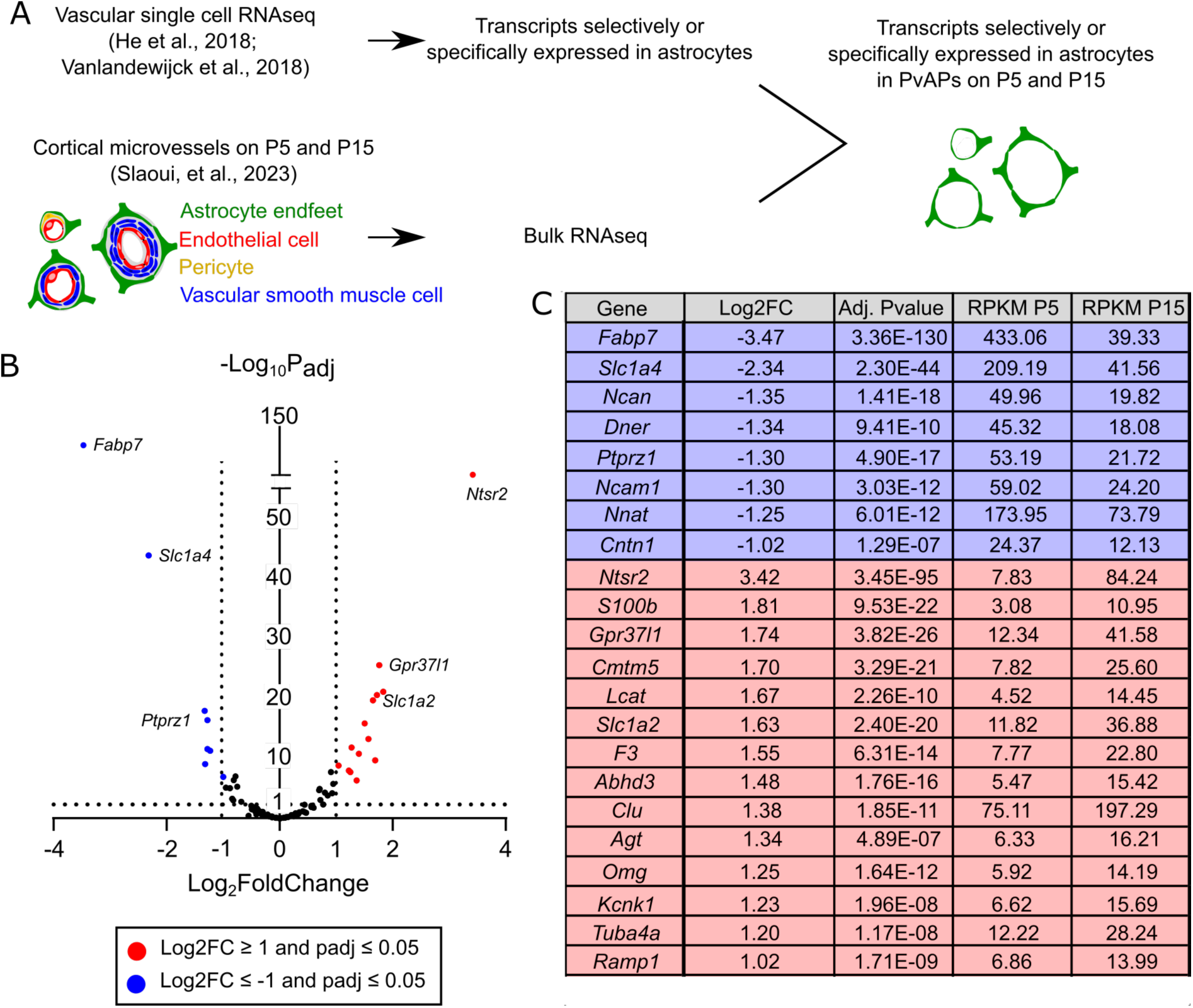
A transcriptomic analysis of mRNAs selectively or specifically expressed in astrocytes and present in PvAPs during postnatal development. **A.** Flowchart for the identification of mRNAs selectively or specifically expressed in astrocytes and present in PvAPs. These transcripts were identified in previously published single-cell RNAseq datasets (He et al., 2016; Vanlandewijck et al., 2018) (**Table S1**) and compared with a previous transcriptional analysis of MVs extracted from the dorsal cortex on P5 and P15 (Slaoui et al., 2023). In the purified MVs, PvAPs (in green) remain attached to the vessel surface, which allows the identification of mRNAs selectively or specifically expressed in astrocytes and present in cortical PvAPs on P5 and P15. **B.** Volcano plot of the astrocytes’ transcriptional changes between P5 and P15 in MVs purified from whole brain. mRNA with similar expression levels on P5 and P15 (-1 < log_2_FC < 1 or padj > 0.05) are shown as black dots; transcripts upregulated on P15 (log_2_FC ≥ 1 and padj ≤ 0.05) are represented as red dots; transcripts downregulated on P15 (logFC ≤ - 1 and padj ≤ 0.05) are represented as blue dots; the dotted lines indicate the threshold for a padj of 0.05 and Log_2_FoldChange of -1 and 1. **C.** List of mRNAs selectively or specifically expressed in astrocytes and up-or downregulated in purified MVs on P15, compared with P5.

To confirm these results and further analyze the developmental profile of mRNAs present in PvAPs, we next purified whole brain MVs on P5, P10, P15, and P60 and performed a qPCR analysis (**Fig. 4 *left columns***). We focused on a selection of mRNAs that were either upregulated on P15 vs. P5 *(Gpr37l1, Slc1a2, Ntsr2*) (**Fig 4A**), unchanged (*Mlc1, Hepacam, Kcnj10, Slc1a3* and *Aqp4*), or downregulated on P15 vs. P5 (*Fabp7, Slc1a4* and *Ptprz1*) (**Fig. 4B-D**). The transcripts found to be down-or upregulated using RNASeq (**Table S1**) had similar qPCR profiles on P5 and P15 (**Fig. 4**); this indicated that RNASeq results in the cortex for these transcripts are representative of the whole brain. In contrast, transcripts not found to be down-or upregulated in the RNASeq study (**Table S1**) were all upregulated in qPCR experiments on P15 whole brain MVs vs. P5, which indicated the presence of regional differences (**Fig. 4**). Interestingly, some mRNAs were further upregulated (*Gpr37l1, Mlc1, Kcnj10, Slc1a2, Ntsr2* and *Aqp4*) or downregulated (*Fabp7, Ptprz1* and *Slc1a4*) between P15 and P60, i.e. after the completion of glial coverage (**Fig. 4**).

**Figure 4.**
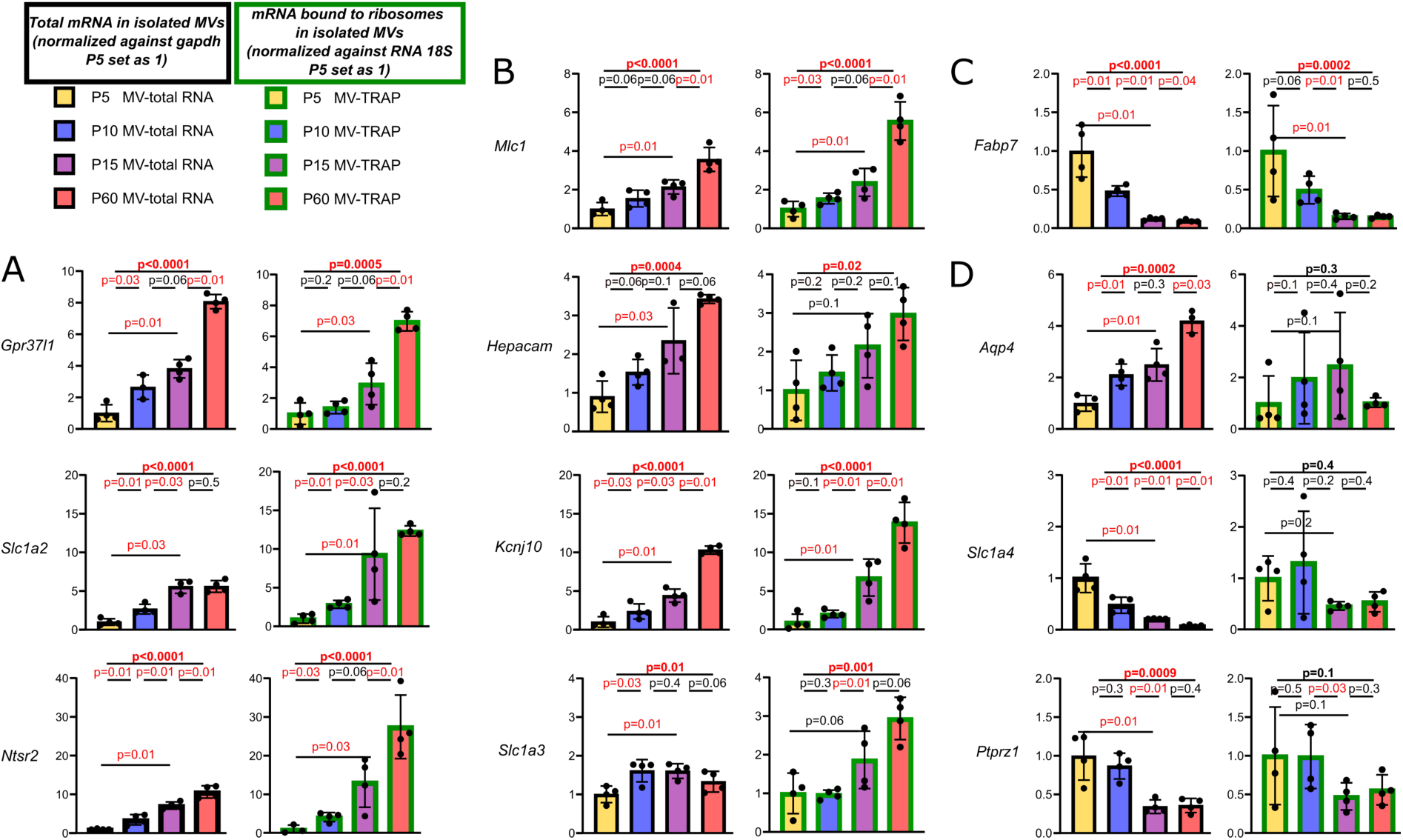
Developmental profile of mRNAs selectively or specifically expressed in astrocytes and polysomes in PvAPs. qPCR analysis of mRNAs selectively or specifically expressed in astrocytes and present in developing PvAPs. The histograms on the left (black-boxed bars) correspond to experiments performed on total mRNAs extracted from whole-brain-purified MVs. The results were normalized against *Gapdh*. The histograms on the right (green-boxed bars) represent experiments performed on mRNAs bound to ribosomes extracted with TRAP from whole-brain Aldh1l1:L10a-eGFP purified MVs. The results were normalized against *RNA 18S*. Levels on P5 are set to 1. Experiments were conducted on P5, P10, P15, and P60. The histograms show the mean ± SD. Kruskal-Wallis test (overall, in bold) and one-tailed Mann-Whitney test (comparison of stages). n=3 to 4 samples per stage (mice per sample: 5 for P5; 3 for P10; 3 for P15; 2 for P30; 2 for P60). Significant p-values (<0.05) are shown in red. The raw data are presented in **Table S2**. Several different profiles were found: a progressive increase in total and polysomal mRNA levels (**A, B**), a progressive decrease in total and polysomal mRNA levels (**C**), and discordant results (an increase in total mRNA levels and a decrease in polysomal mRNA levels, or vice versa) (**D**).

We next asked whether the distribution of PvAP mRNAs might be altered in an animal model of MLC, in which PvAP development is impaired (Gilbert et al., 2021). This rare leukodystrophy is primarily caused by the absence of the PvAP transmembrane protein MLC1 (Leegwater et al., 2001; Topcu et al., 2000). MLC patients display early-onset macrocephaly and progressive white matter vacuolation, which leads to slowly progressing ataxia, spasticity, and cognitive decline (Duarri et al., 2008; Lanciotti et al., 2016; Leegwater et al., 2001). The absence of MLC1 alters the astrocytes’ morphological maturation, polarity and perivascular organization from P10 onwards, with discontinuous PvAPs and the loss of PvAP polarity and cohesiveness (Gilbert et al., 2021). PvAP mRNAs cannot be extracted from *Mlc1* KO mice because the PvAPs are poorly cohesive and are lost during MV purification (Gilbert et al., 2021). Thus, we used FISH to analyze mRNA distribution on P15 sections of the somatosensory cortex in *Mlc1* KO mice and WT littermates (**Fig. 5A**). We focused on *Gpr37l1*, the gene coding for a putative prosaposin receptor (Smith, 2015). Gpr37l1 is a ligand of MLC1 and has been shown to regulate MLC1 targeting to the membrane in primary astrocytes and in the Bergmann glia of the cerebellum (Alonso-Gardon et al., 2021). It was recently reported that, like MLC1, Gpr37l1 regulates the morphological maturation of astrocytes (Nguyen et al., 2023). By staining blood vessels for IB4 and quantifying the FISH dots in the parenchyma and at the vessel surface, we found that *Mlc1* KO displayed lower levels of *Gpr37l1* mRNAs in both whole astrocytes and in PvAPs compared to WT **(Fig. 5A, B)**.

**Figure 5.**
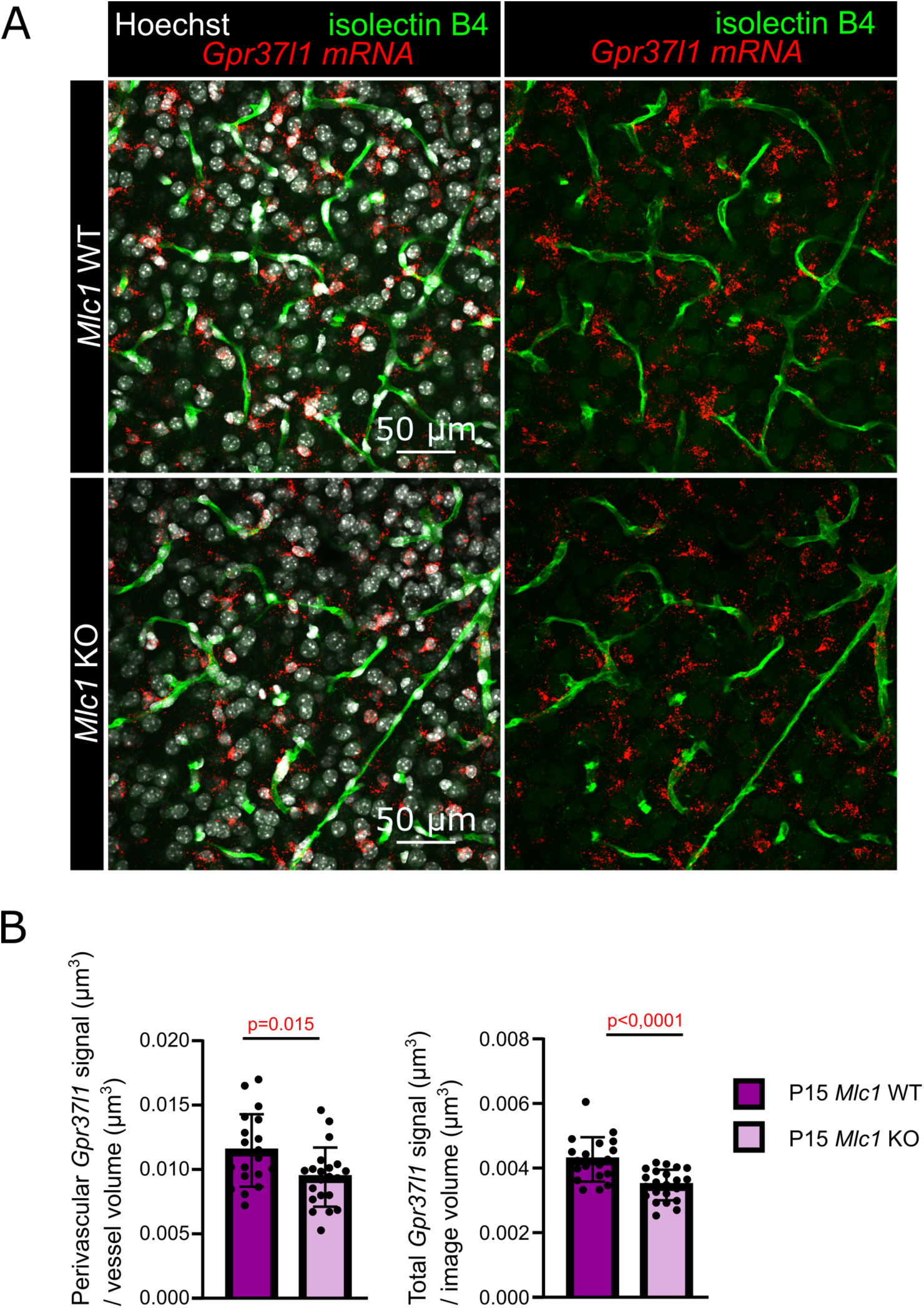
Perivascular distribution of *Gpr37l1* mRNA in *Mlc1* KO astrocytes. **A.** Representative confocal microscopy images of FISH detection of *Gpr37l1* mRNAs in the P15 somatosensory cortex of *Mlc1* KO mice or WT littermates. Blood vessels (in green) were stained with isolectin B4. **B.** Quantification of the perivascular FISH signal normalized against the vessel volume (left) and the total FISH signal normalized against the image volume (right), under each condition. The data are presented as the mean ± SD (*Mlc1* KO: n=21 images in 4 mice; WT: n=19 images in 4 mice); unpaired t-tests (two-tailed). The raw data are presented in **Table S2**.

In summary, we identified a pool of astrocyte-selective or -specific transcripts present in PvAPs and determined their postnatal developmental profile. We found that the levels of various mRNAs change from P5 to P60, i.e. during and after the formation of PvAPs. We also found that the P15 distribution of *Gpr37l1* mRNA in cortical PvAPs is altered in a mouse model of MLC.

### Postnatal developmental profiles of polysomal mRNAs in PvAPs

We next looked at the translational status of PvAP mRNAs during postnatal development. We used our refined translation affinity purification (TRAP) protocol to extract PvAP polysomal mRNAs from MVs purified from Aldh1l1:L10a-eGFP mouse whole brain on P5, P10, P15, and P60. In qPCR assays, most of polysomal mRNAs showed the same developmental changes as total mRNA did (**Fig. 4 right columns**). Amounts of *Gpr37l1, Slc1a2, Ntsr2, Mlc1, Hepacam, Kcnj10* and *Slc1a3* mRNAs increased from P5 to P60 (**Fig. 4A and B**). The increase in the amount of polysomal *Slc1a3* mRNA was delayed, relative to the increase in total mRNA (**Fig. 4B**). The amount of *Fabp7* decreased from P5 to P60 (**Fig. 4C**). However, discordant developmental profiles were observed when comparing *Aqp4, Slc1a4* and *Ptprz1* polysomal and total mRNAs (**Fig. 4D**).

These results suggested that distribution and translation of specific mRNAs occur in PvAPs during postnatal development. The developmental profiles of total mRNAs and polysomal mRNAs in PvAPs were generally correlated, which suggests that mRNA transport has a crucial role in the organization of local translation. They can also differ, which suggests that local translation in PvAPs is regulated at several different levels.

### Postnatal developmental profiles of proteins in PvAPs

Given that translation rates and protein levels are not necessarily correlated, we measured the levels of some PvAP proteins on Western blot of proteins extracted from P5 to P60 purified MVs (i.e. with attached PvAPs) from whole brain (**Fig. 6A**). Fabp7, Glt1 (encoded by *Slc1a2*), Kir4.1 (encoded by *Kcnj10*) had the same developmental profiles as polysomal mRNAs (**Fig. 6B**). In contrast, Aqp4 levels increased significantly from P5 to P60, while polysomal mRNAs did not change significantly during this time.

**Figure 6.**
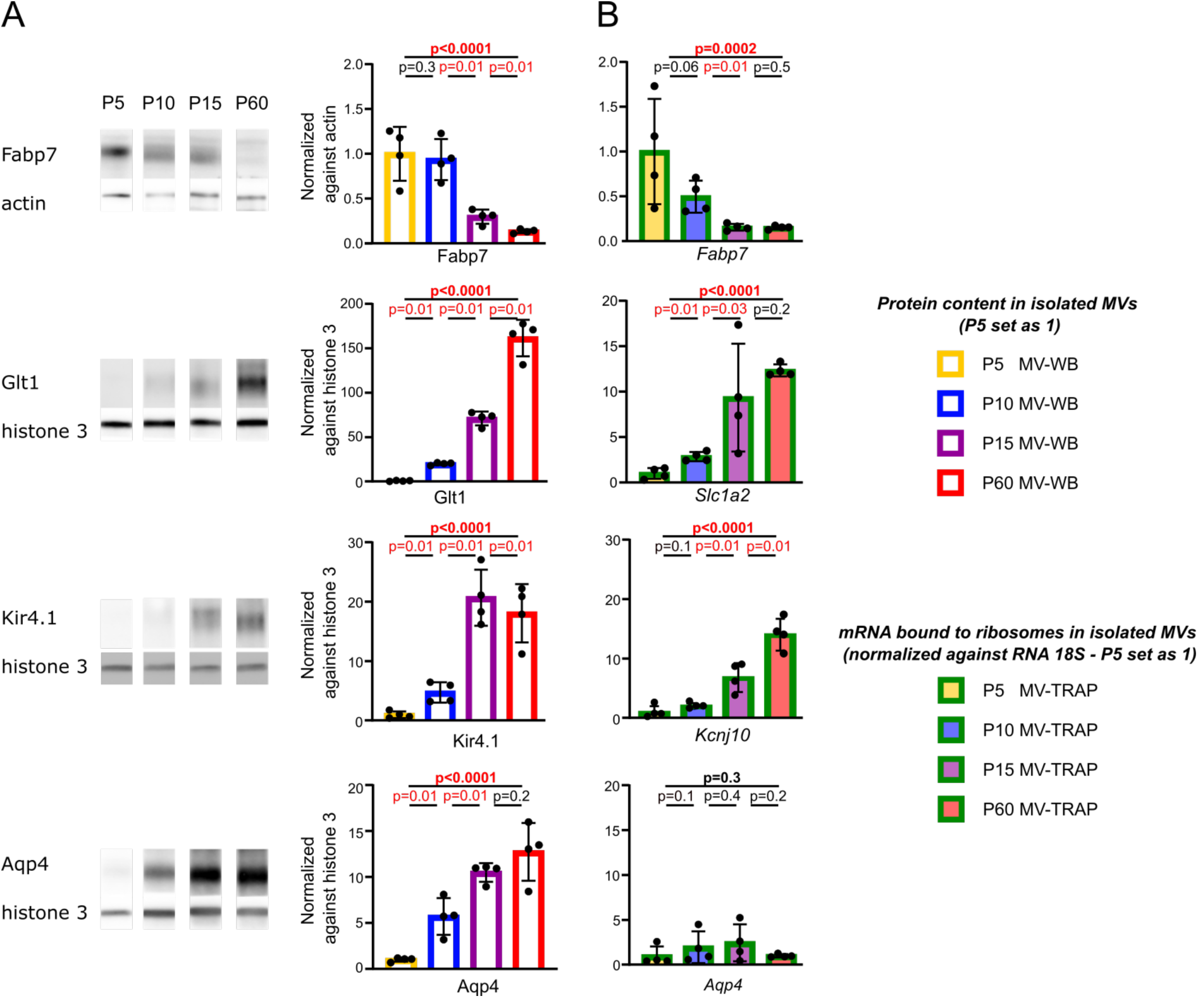
Comparisons of local translation and protein levels in developing PvAPs. **A.** Western blot detection, analysis, and quantification of proteins selectively or specifically expressed in astrocytes and present in extracts of MVs purified from whole brains (with attached PvAPs) on P5, P10, P15, and P60. The signals were normalized against actin or histone 3. P5 levels were set to 1. The histograms show the mean ± SD (n=4 samples per developmental stage; mice per sample: 5 for P5; 4 for P10; 3 for P15; 2 for P60); Kruskal-Wallis test (overall, in bold) and one-tailed Mann-Whitney test (comparison of stages). The raw data are presented in **Table S2**. **B.** Data on the qPCR detection of mRNA bound to ribosomes and extracted with TRAP from whole brain Aldh1l1:L10a-eGFP purified MVs on P5, P10, P15, and P60 (**see** Fig. 4 **right columns**). The results were normalized against *RNA 18S*. P5 levels were set to 1. Experiments were conducted on P5, P10, P15, and P60. The histograms show the mean ± SD. Kruskal-Wallis test (overall, in bold) and one-tailed Mann-Whitney test (comparison of stages). Significant p-values (<0.05) are shown in red. n=4 samples per stage (mice per sample: 5 for P5; 3 for P10; 3 for P15; 2 for P60). The raw data are presented in **Table S2**.

The correlation between polysomal mRNA levels and protein levels in PvAPs suggests that local translation contributes to the postnatal molecular maturation of PvAPs. However, the fact that some results were not correlated suggests that other mechanisms influence the protein repertoire of PvAPs during postnatal development.

### Postnatal developmental profiles of polysomal mRNAs in PvAPs and astrocytes

We have previously reported that in the adult brain, some polysomal mRNAs are highly enriched in PvAPs (relative to the soma); this enrichment strongly suggests that local translation sustains the astrocyte’s perivascular polarity (Boulay et al., 2017). Hence, we next sought to determine whether this polarity is established during postnatal development. Using Aldh1l1:L10a-eGFP mice, we extracted polysomal mRNAs (i) from PvAPs by performing TRAP on purified MVs, and (ii) from whole astrocytes by performing TRAP on whole brain on P5, P10, P15, and P60. The mRNAs were quantified by qPCRs, and the MV/whole brain ratio (considered as a polarity index) was calculated for each time point (**Fig. 7**). For *Mlc1*, *Slc1a2*, *Kcnj10*, *Ntsr2*, *Slc1a3*, *Hepacam, Ptprz1* and *Slc1a4,* the MV/whole brain ratio did not change between P5 and P15 but was higher on P60 - suggesting that these polysomal mRNAs are highly polarized towards PvAPs from P15 onwards. In contrast, no change was observed for *Aqp4, Fabp7* and *Gpr37l1*, which indicates that, for these genes, the enrichment of polysomal mRNAs in PvAPs does not change postnatally.

**Figure 7.**
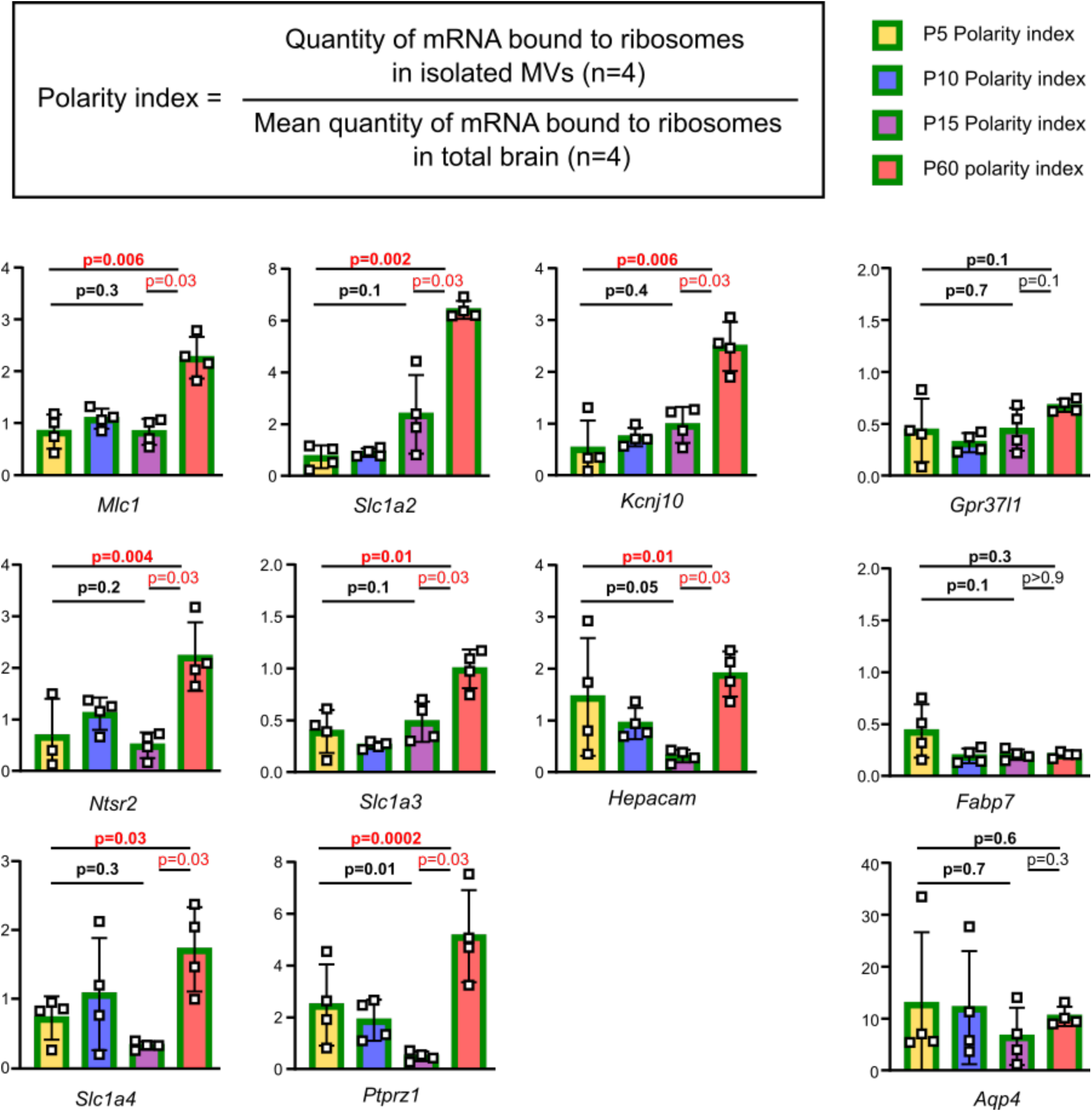
Postnatal developmental profiles of polysomal mRNA in PvAPs and astrocytes. The ratio for qPCR levels of polysomal mRNAs extracted with TRAP on Aldh1l1:L10a-eGFP MVs from whole brain on P5, P10, P15, and P60. The histograms show the mean ± SD. Kruskal-Wallis tests (overall and P5-P15, in bold) and two-tailed Mann-Whitney test (P15-P60). n= 3 to 4 samples per stage (mice per MV sample: 5 for P5; 3 for P10; 3 for P15; 2 for P60; 1 mouse per sample for the whole-brain analysis). Significant p-values (<0.05) are shown in red. The raw data are presented in **Table S2**.

Taken as a whole, these results suggest that the distribution of specific polysomal mRNAs is highly polarized in PvAPs from P15 onwards.

## Discussion

In the brain, the translation of distally localized mRNAs is crucial for neuronal development; it contributes to axon growth, branching and synaptogenesis (for a review, see (Agrawal and Welshhans, 2021)). The transport and translation of ß-actin-encoding mRNAs (shown to sustain dendritic arborization and axon growth) have been extensively studied (Jung et al., 2014; Perycz et al., 2011; Wong et al., 2017). More generally, the axon growth cone transcriptome of an intracortical projection in the early postnatal mouse brain was recently described (Engmann et al., 2022; Poulopoulos et al., 2019). Shigeoka et al. have sequenced the translatome from developing and adult retinotectal axons (Shigeoka et al., 2016). Local translation has also been studied in radial glial cells, i.e. proliferative neural stem cells (Pilaz et al., 2016). mRNA trafficking and local translation might be critical for the production of progenitor cells, cytoskeletal dynamics, and thus the regulation of cortical development (D’Arcy and Silver, 2020). It was recently shown that the local sorting and local translation of mRNA encoding the GTPase-activating protein ARHGAP11A in the basal endfeet of radial glial cells controlled the latter’s morphology and mediated neuronal positioning (Pilaz et al., 2023). Considering the importance of dynamic mRNA distribution and local translation in the development of the brain, we hypothesized that these mechanisms are involved in astrocyte development. In the mouse, astrocytes are produced from radial glial cells during the first postnatal week. The proliferation and spatial dispersion of newly differentiated astrocyte progenitors in the cortex are followed by a maturation phase, with an increase in volume and morphological complexity between P7 and P21 (Clavreul et al., 2019). Astrocytes probably acquire their final subtype characteristics by interacting with their environment in general and neurons and brain vessels in particular. In recent research, we showed that cortical and hippocampal perivascular astrocytic coverage develops around the time of birth, doubles between P5 and P10, and is complete on P15 (Gilbert et al., 2021). Interestingly, this time window corresponds to a molecular maturation phase during which PvAPs acquire specific molecular properties, such as the MLC1/GlialCAM complex (Gilbert et al., 2021; Lunde et al., 2015). However, the underlying mechanisms of PvAP development and molecular maturation are poorly understood.

By combining *in situ* and molecular analyses in the present study, we found that translation is already present in newly formed PvAPs on P5, with ribosomes, mRNAs, polysomes and puromycilated nascent protein chains. The perivascular volume occupied by astrocytic eGFP-tagged ribosomes increased progressively after birth and paralleled the development of PvAPs (as quantified by the amount of perivascular eGFP in Aldh1L1:eGFP mice). Our subcellular observations revealed that about half of the PvAPs are equipped with RER or free polysomes on P5. This proportion did not change significantly between P5 to P60. Hence, these results indicate that the formation of PvAPs is accompanied by mRNA and polysome transport, and extension of the RER network. We observed a Golgi apparatus in some PvAPs on P5, which suggests that protein maturation and secretion pathways are present in developing PvAPs. It has been reported that about 7% of mature PvAPs in the cortex and hippocampus are equipped with a Golgi apparatus (Boulay et al., 2017). The observation that not all PvAPs are equipped with RER, free polysomes and/or a Golgi apparatus indicates that local translation does not occur in all PvAPs and that locally translated proteins might not follow the same post-translational modifications in all PvAPs; this adds a significant level of complexity to the PvAPs’ molecular repertoire and functions.

Total mRNAs and polysomal mRNAs have already been observed in adult PvAPs from whole-brain preparations (Boulay et al., 2017). Here, we hypothesized that dynamic mRNA distribution and local translation could sustain the postnatal molecular maturation of PvAPs. To address this hypothesis, we focused on mRNAs that are selectively or specifically expressed in astrocytes, according to published single-cell RNASeq datasets (He et al., 2016; Vanlandewijck et al., 2018). Using previously published transcriptomic data, we then analyzed the mRNAs’ expression in cortical MVs purified on P5 and P15, (Boulay et al., 2015b; Slaoui et al., 2023). During the mechanical purification of MVs, PvAPs remain attached to the vessel surface and astrocyte somas are lost (Boulay et al., 2017; Boulay et al., 2015b). Thus, astrocyte-selective or -specific transcripts in purified MVs necessarily come from PvAPs. Our results showed that not all astrocyte mRNAs are present in PvAPs; hence, mRNA transport probably sustains the establishment of a specific molecular identity in developing PvAPs. Furthermore, our results indicate that astrocytic mRNAs present in PvAPs have different developmental profiles. The increase in the level of some mRNAs between P5 and P10 might be related to the astrocytic perivascular coverage, which is much higher on P10 than on P5 (Gilbert et al., 2021). However, by focusing on a selected set of mRNAs, we showed that levels of *Gpr37l1, Mlc1, Kcnj10,* and *Ntrs2* mRNAs continue to rise after P15 (when astrocytic coverage is almost complete) and that levels of *Fabp7* and *Slc1a4* mRNAs fell progressively from P5 to P60. Thus, mRNA levels might be specifically controlled during the postnatal development and maturation of PvAPs.

As it has already been described for axon growth cones, subtranscriptomes and subproteomes are not necessarily correlated (Poulopoulos et al., 2019). To further characterize local translation in PvAPs, we compared (i) levels of total mRNA with levels of polysomal mRNAs (extracted with TRAP), and (ii) levels of polysomal mRNA with protein levels. For most transcripts, the changes over time in total mRNAs, polysomal mRNAs and proteins were similar; this indicates that the transport of mRNAs into PvAPs fueled local translation. However, some discrepancies were also observed. In particular, levels of total *Aqp4* mRNA and Aqp4 protein rose progressively from P5 to P60, while polysomal *Aqp4* mRNA levels were similar at the two time points (albeit with a great level of variability during that time period). These observations suggest that a proportion of the *Aqp4* mRNA transported to PvAPs during postnatal development is not bound to ribosomes and is in a translationally repressed state. Considering that mRNA transport requires high levels of energy, this pool of mRNAs might serve as a local supply and might be reactivated under specific circumstances. The constant increase in the level of Aqp4 protein in PvAPs (in contrast to levels of polysomal mRNAs) also suggests that protein-level regulation (such as membrane stabilization and transport from the soma) also influences the final quantity of Aqp4 in PvAPs. Taken as a whole, these results indicate that local translation in PvAPs is an important mechanism for the postnatal acquisition of the PvAP’s molecular identity. The fact that levels of total mRNAs, polysomal mRNAs and proteins were not necessarily correlated in PvAPs strongly suggests that the mechanisms regulating the fate of each mRNA also contribute actively to the developmental process.

Local translation is altered in several neurodegenerative diseases, including amyotrophic lateral sclerosis (Piol et al., 2023) and Alzheimer’s disease (Baleriola and Hengst, 2015; Baleriola et al., 2014). Since local translation is required during brain development, its impairment might be linked to neurodevelopmental diseases - as has been suggested for autism (Chen et al., 2019). It has been shown that fragile X messenger ribonucleoprotein (FMRP) regulates the transport and translation of a local transcriptome in radial glial cells (Pilaz et al., 2016). Here, we showed that the expression and distribution of *Gpr37l1* RNAs in PvAPs were altered in the *Mlc1* KO mouse model of MLC (Hoegg-Beiler et al., 2014). Interestingly, MLC1 and Gpr37l1 interact in astrocytes (Alonso-Gardon et al., 2021). Furthermore, the absence of MLC1 or Gpr37l1 has been shown to impair the morphological maturation of astrocytes (Gilbert et al., 2021; Nguyen et al., 2023). The absence of MLC1 also alters the course of astrocytic perivascular coverage (Gilbert et al., 2021). Taken as a whole, these results suggest that MLC1 may influence astrocyte morphology by controlling the expression and distribution of *Gpr37l1* mRNA in astrocytes.

In summary, we found that newly formed PvAPs are equipped with translation machinery and perform local translation. We also demonstrated that the levels of astrocytic mRNAs and polysomal mRNAs in PvAPs change between P5 and P60. Lastly, we showed that mRNA distribution in PvAPs is modified in a mouse model of MLC. We therefore suggest that dynamic mRNA distribution and local translation contribute to the postnatal maturation of PvAPs and, more generally, to the development of the gliovascular unit.

## Materials and Methods

### Mice

Tg(Aldh1l1-eGFP/Rpl10a) JD130Htz (MGI: 5496674) (Aldhl1:L10a-eGFP) and Tg(Aldh1l1-eGFP) OFC789Gsat/Mmucd (Aldh1l1:eGFP) mice were obtained from Nathaniel Heintz (Rockefeller University, NY) and kept under pathogen free conditions. These mouse strains have been generated by bacterial artificial chromosome transgenesis (Doyle et al., 2008; Gong et al., 2003; Heiman et al., 2008). The Aldh1l1 promoter is used to induce expression of L10a-eGFP or eGFP in astrocytes and is also active in neurogenic areas (Foo and Dougherty, 2013). Western blot experiments were performed on Swiss mice. *Mlc1* KO mice were provided by Raul Estevez (Barcelona, Sapin) and maintained on a C57BL6 genetic background (Hoegg-Beiler et al., 2014). The experiments and techniques reported here complied with the ethical rules of the French national agency for animal experimentation.

### Microvessel purification

Microvessels with associated PvAPs were purified from whole brain, as described previously (Boulay et al., 2015b). Briefly, brains were resuspended in HBSS/HEPES, using an automated Dounce homogenizer. After an initial centrifugation at 2000 g for 10 min, the pellet was resuspended in HBSS/Dextran 18% and centrifuged at 4000 g for 15 min to separate the myelin from the vessels. This new pellet contained the brain vessels with attached PvAPs and was then resuspended in HBSS/BSA 1%. Filtration through a 100 µm mesh filter removed large arteries and veins. The eluate was then filtered on a 20 µm mesh filter. The retained MVs with attached PvAPs consisted of arterioles, venules, and capillaries.

### RNA sequencing and analysis

RNASeq data were analyzed as described previously (Slaoui et al., 2023). The RNASeq gene expression data and the raw fastq files are available on the GEO repository (www.ncbi.nlm.nih.gov/geo/), with the accession number GSE173844.

### Identification of astrocyte-specific or -selective transcripts

Raw reads from GEO datasets GSE99058 and GSE98816 were downloaded (He et al., 2016; Vanlandewijck et al., 2018). Seurat 3.1.1 was used to normalize the unique molecular identifiers (Butler et al., 2018), using a global-scaling method, a scale factor of 10000, and log-transformation of the data. This was followed by a linear transformation scaling step, so that highly expressed genes did not have an excessively high weighting in the downstream analysis. A transcript was considered as to be specific to or preferentially expressed in astrocytes when it was detected in more than 60% of astrocytes, detected in less than 12% cells of other cell types, and had a significant higher expression level in astrocytes than in other cells (according to a Wilcoxon rank sum test, and with log_2_FC >1.5) (**Table S1**). The expression level of each transcript in MVs on P5 and P15 was assessed in RPKM.

### Transmission electron microscopy

Mice were anesthetized with ketamine-xylazine (140 and 8 mg/kg, respectively, i.p.) and transcardially perfused with the fixative (2% paraformaldehyde, 3% glutaraldehyde, and 3 mM CaCl_2_ in 0.1M cacodylate buffer pH 7.4) for 12 min. The brains were removed and left overnight at 4 °C in the same fixative. Brain fragments (0.3 x 1 x 1 mm^3^) were postfixed first in 0.1M cacodylate buffer pH 7.4 + 1% OsO_4_ for 1h at 4 °C and then in 1% aqueous uranyl acetate for 2 h at room temperature (RT). After dehydration in graded ethanol and then propylene oxide, the fragments were embedded in EPON^TM^ epoxy resin (Electron Microscopy Sciences, Hatfield, PA). Ultrathin (80 nm) sections were prepared, stained with lead citrate, and imaged in a Jeol 100S transmission electron microscope (Jeol, Croissy-sur-Seine, France) equipped with a 2k x 2k Orius 830 CCD camera (Roper Scientific, Evry, France).

### Translating ribosome affinity purification (TRAP)

Whole brain or MV homogenates from Aldh1l1:L10a-eGFP mice underwent TRAP by immunoprecipitating eGFP-fused astrocytic polyribosomes with anti-GFP antibodies and protein-G-coupled magnetic beads, as described elsewhere (Boulay et al., 2019).

### Quantitative RT-PCR

RNA was extracted from MV homogenates or TRAP extracts using the Rneasy Lipid Tissue Mini Kit (Qiagen, Hilden, Germany). cDNA was then generated using the Superscript™ III Reverse Transcriptase Kit (ThermoFisher). Differential cDNA expression was measured using droplet digital PCR. Briefly, cDNA and primers (**Table S3**) were distributed into approximately 10,000 to 20,000 droplets. cDNAs were then PCR-amplified in a thermal cycler and read (as the numbers of positive and negative droplets) with a QX200 Droplet Digital PCR System (Bio-Rad). The ratio for each tested gene was normalized against the number of droplets positive for *RNA 18S* (in the case of polysomal mRNAs extracted with TRAP) or for *Gapdh* (in the case of total mRNA).

### Western blots

Purified MVs with attached PvAPs were resuspended in 50 µL of SDS 2%, sonicated three times for 10 s at 20 Hz (Vibra cell VCX130), and boiled in Laemmli loading buffer. Protein content was measured using the Pierce 660 nm protein assay reagent (Thermo Scientific, Waltham, MA, USA). Equal amounts of proteins were separated by denaturing electrophoresis in Mini-Protean TGX stain-free gels (Bio-Rad) and then electrotransferred to nitrocellulose membranes using the Trans-blot Turbo Transfer System (Bio-Rad). Membranes were hybridized as described previously (Ezan et al., 2012). The antibodies used in the present study are listed in **Table S3**. Horseradish peroxidase activity was visualized using an enhanced chemiluminescence assay in a Western Lightning Plus system (Perkin Elmer, Waltham, MA, USA). Chemiluminescent imaging was performed on a FUSION FX system (Vilber, South Korea). Four independent samples were analyzed in each experiment. The level of chemiluminescence for each antibody was normalized against the signal for actin or histone 3.

### Immunohistochemical analysis

Mice were anesthetized with pentobarbital (600 mg/kg, i.p.) and killed by transcardiac perfusion with PBS/PFA 4%. The brain was removed and cut into 40-µm-thick sections using a Leitz (1400) cryomicrotome. Brain slices on glass slides were immersed in the blocking solution (PBS/normal goat serum 5%/Triton X-100 0.5%) for 1 h at RT and then incubated with primary antibodies (**Table S3**) diluted in the blocking solution 12 h at 4 °C. After 3 washes in PBS, slices were incubated for 2 h at RT with secondary antibodies and Hoechst reagent, rinsed in PBS, and mounted in Fluoromount G (Southern Biotech, Birmingham, AL) for confocal analysis. Tissues were imaged using a 40X objective on a Zeiss Axio-observer Z1 with a motorized XYZ stage (Zeiss, Oberkochen, Germany).

### High-resolution fluorescent *in situ* hybridization

Fluorescent *in situ* hybridization (FISH) was performed on floating PBS/PFA 4% fixed brain sections, according to the v2 Multiplex RNAscope technique (Advanced Cell Diagnostics, Inc., Newark, CA, USA) described previously (Oudart et al., 2020). Brain sections were treated with protease at RT. After the FISH, brain sections were stained with isolectin B4 (1/100) in PBS/normal goat serum 5%/Triton 0.5% overnight at 4°C. Nuclei were stained with Hoechst reagent (1/2000). The brain sections were imaged using a Spinning Disk CSU-W1 microscope and Metamorph Premier 7.8 software.

### FISH and immunofluorescence quantification

The data were analyzed using a custom-written plug-in (https://github.com/orion-cirb/Vessel_IB4/) developed for Fiji software (Schindelin et al., 2012), using Bio-Formats (Linkert et al., 2010), CLIJ (Haase et al., 2020), GDSC (https://github.com/aherbert/gdsc) and 3D Image Suite libraries (Ollion et al., 2013) . In brief, the vessel channel was first filtered with a two-dimensional (2D) Laplacian of Gaussian filter (σ = 20) and segmented with the triangle method. The binary mask thus obtained was labeled in three dimensions, and three-dimensional (3D) vessels were filtered by volume to avoid false positives. In a second step, RNA FISH dots were detected by successively applying a 3D difference of Gaussians filter (σ_1_ = 1, σ_2_ = 2) and the moments thresholding method. Dots were then labeled in 3D, filtered by volume, and classified as perivascular or not as a function of their distance to the closest vessel: dots with a distance below 2 µm were considered to be perivascular. For each category, the number of dots, the dots’ volume, and their integrated intensity were computed.

Another custom plug-in was written for immunofluorescence (https://github.com/orion-cirb/Astrocytes_InOut_Vessels). Vessels were detected by filtering the channel with a 2D Laplacian of Gaussian filter (σ = 14) and segmented with the triangle method. Labeling the segmentation yielded 3D vessels by volume. eGFP and L10a-eGFP signals were detected by applying first a 2D median filter (radius = 2) and then an automatic thresholding method. The mean and Huang methods were used for L10a-eGFP and eGFP staining, respectively. Lastly, the volume of the contact between L10a-eGFPand eGFP signals and vessels was computed.

### Puromycilation assay

P5 Aldh1l1:eGFP mice were anesthetized with ketamine-xylazine (140 and 8 mg/kg, respectively, i.p.) and injected intraperitoneally with 50 µL of puromycin (P7255, Sigma Aldrich) diluted in water (10 mg/mL; 225mg/kg). The same volume of water was injected into control mice. After 25 min., mice were killed by transcardiac perfusion with PBS/PFA 4%, and brain sections were prepared. Puromycin was detected by immunohistochemistry, as described above (**Table S3)**.

## Acknowledgements

We thank Raul Estevez for the kind gift of *Mlc1* KO mice.

